# Modelling dysfunction-specific interventions for seizure termination in epilepsy

**DOI:** 10.1101/2025.02.24.639968

**Authors:** Aravind Kumar Kamaraj, Matthew Parker Szuromi

## Abstract

Epileptic seizures result from abnormal synchronous neuronal firing caused by an imbalance between excitatory and inhibitory neurotransmission. While most seizures are self-limiting, those lasting over five minutes, termed status epilepticus, require medical intervention. Benzodiazepines, the first-line treatment, terminate seizures by enhancing GABAergic inhibition, but fail in approximately 36% of cases. In this paper, we employ a neural mass framework to investigate how different interventions influence brain dynamics and facilitate seizure termination. As seizures are characterized by persistent firing, we extend the classic Wilson-Cowan framework by introducing a term called sustenance which encodes factors that promote or discourage perpetual firing. The resulting model captures transitions between normal activity and seizure and provides a tractable framework for analysing diverse pathophysiological mechanisms. We first show how various dysfunctions — such as hyperexcitation, depletion of inhibitory neurotransmitters, and depolarizing GABAergic transmission — can all give rise to seizures, with overlapping but distinct dynamics. Building on this foundation, we turn to the central question of intervention: how different treatments act on these mechanisms to terminate seizures. We find that while enhancing GABAergic inhibition is generally effective, it fails when GABA becomes depolarizing. In such cases, interventions like levetiracetam that suppress sustained excitatory activity remain effective. These findings highlight the importance of aligning interventions to the specific underlying dysfunction for effective seizure termination.

## I. INTRODUCTION

Epilepsy is a neurological disorder characterized by recurrent, unprovoked seizures caused by abnormal brain activity. Seizures are episodes of abnormal, excessive, and synchronous neuronal activity in the brain that can lead to transient disruptions in behaviour, sensation, or consciousness. Affecting over 50 million people globally, epilepsy is among the most common neurological conditions [1]. Disruptions in the balance between excitatory and inhibitory neurotransmission play a key role in its pathophysiology; however, the precise mechanisms by which epileptic seizures begin and end are not well understood [2]. Experimental studies have demonstrated that both enhanced excitation [3, 4] and impaired inhibition [5–7] can lead to seizures. Further, as seizure onset, prediction, and prevention carry greater immediate clinical relevance, they have received considerable research attention, whereas the mechanisms underlying seizure termination remain comparatively under-explored [8].

Medications are required to terminate a seizure when it becomes prolonged or fails to resolve spontaneously. While most seizures are self-limiting, those lasting over five minutes, termed *status epilepticus*, pose a high risk of neuronal injury, systemic complications, and increased mortality [9]. Benzodiazepines are the first-line treatment due to their rapid onset and potent anticonvulsant effects [10]. They terminate seizures by enhancing GABA-mediated inhibition [11]. Gamma-aminobutyric acid (GABA), the primary inhibitory neurotransmitter in the mature central nervous system, exerts its effects primarily through GABA_*A*_ receptors on neurons [12]. Activation of these receptors opens chloride channels, resulting in chloride influx and neuronal hyperpolarization, thereby reducing neuronal excitability [13].

However, approximately 36% of status epilepticus cases are refractory to benzodiazepine treatment [14]. In such cases, second-line treatments such as levetiracetam, phenytoin, and sodium valproate are recommended [15]. Most of these agents selectively antagonize rhythmic firing caused by excessive excitatory feedback, while sparing normal electrophysiological function [16]. For instance, phenytoin obstructs pathological excitatory feedback through a voltage-dependent blockade of sodium channels responsible for action potential generation [17], whereas levetiracetam works by binding to the SV2A ligand and suppressing the release of the excitatory neurotransmitter glutamate [18, 19].

Given that first-line treatments enhance inhibition while second-line treatments suppress excessive excitation, we hypothesize that aligning the therapeutic strategy with the underlying dysfunction could lead to more effective seizure control. To investigate this, we employ a neural mass model to study how different dysfunctions and pharmacological interventions shape seizure dynamics, offering insights into dysfunction-specific treatment approaches.

Mathematical modelling of epilepsy is a broad field, with models ranging from detailed single-neuron representations to large-scale networks encompassing multiple interconnected brain regions [20]. Neural mass models, situated at the mesoscopic scale, provide a useful level of abstraction for studying the dynamics of a small number of interacting neuronal populations [21, 22]. Earlier models primarily focussed on replicating electroencephalographic (EEG) features during seizures, normal activity, and the transition between the two states [23, 24]. Extensions, such as incorporating distinct fast and slow inhibitory feedback loops, enable the simulation of high-frequency oscillations observed in intracranial EEG recordings [25] while neural field models incorporating a network of neural masses allow for detailed spatiotemporal description of seizure propagation and termination [26, 27].

Additionally, neural mass models have been tailored to study specific epileptic syndromes by including multiple populations, such as thalamic and cortical populations for absence seizures [28], and excitatory subpopulations incorporating depolarizing GABAergic neurotransmission for Dravet’s syndrome [29]. However, as these models evolved to capture increasingly complex dynamics, they have often sacrificed clear physiological interpretation for sophistication, making it challenging to directly map the model parameters and insights to underlying mechanisms.

In this paper, we build upon the original Wilson–Cowan model [30], focusing on a single excitatory and inhibitory population. To capture the defining feature of seizures — persistent neuronal firing — we introduce a term called sustenance, which quantifies factors that either promote or inhibit sustained activity. This term allows us to model both dysfunctions that facilitate seizure-like dynamics and interventions that counteract them. We define the model’s equilibria in physiologically grounded terms: the equilibrium with maximal excitatory activity corresponds to a seizure, and the equilibrium with minimal, non-zero neuronal activity represents normal brain function. While our abstraction does not capture detailed EEG features, it provides an elegant and tractable framework for examining different seizure mechanisms, highlighting the novelty of our approach.

Within this framework, we first investigate how various dysfunctions, such as hyperexcitation, depletion of inhibitory neurotransmitters, and depolarizing GABAergic neurotransmission, can lead to seizure onset. We then turn to the central question of intervention and explore how different treatment strategies act on these mechanisms to terminate seizures.

## II. RESULTS

The dynamical landscape of our model is defined by two first-order ordinary differential equations: one governing excitatory activity, *E*, and the other governing inhibitory activity, *I*. This two-dimensional framework lends itself to a visual and intuitive understanding of brain dynamics, as perturbations to excitatory or inhibitory mechanisms manifest directly as geometric changes in the *E*- and *I*-nullclines. Accordingly, we use phase portraits such as the one shown in Fig. 1, to illustrate these changes and extract qualitative insights. The geometry of the nullclines is shaped by the sigmoidal activation functions, each of which can be divided into three distinct regions: the upper asymptote (*A*_*E*_,*→ A*_*I*_ 1), the lower asymptote (*A*_*E*_, *→A*_*I*_ 0), and the transition curve (the remaining segment in the middle) as shown in the inset in Fig. 1. These three regions correspond to distinct segments of the *E*- and *I*-nullclines, which we use to interpret the model’s dynamics. We begin our analysis by identifying the equilibria that emerge at the intersections between these nullcline segments.

**FIG. 1.**
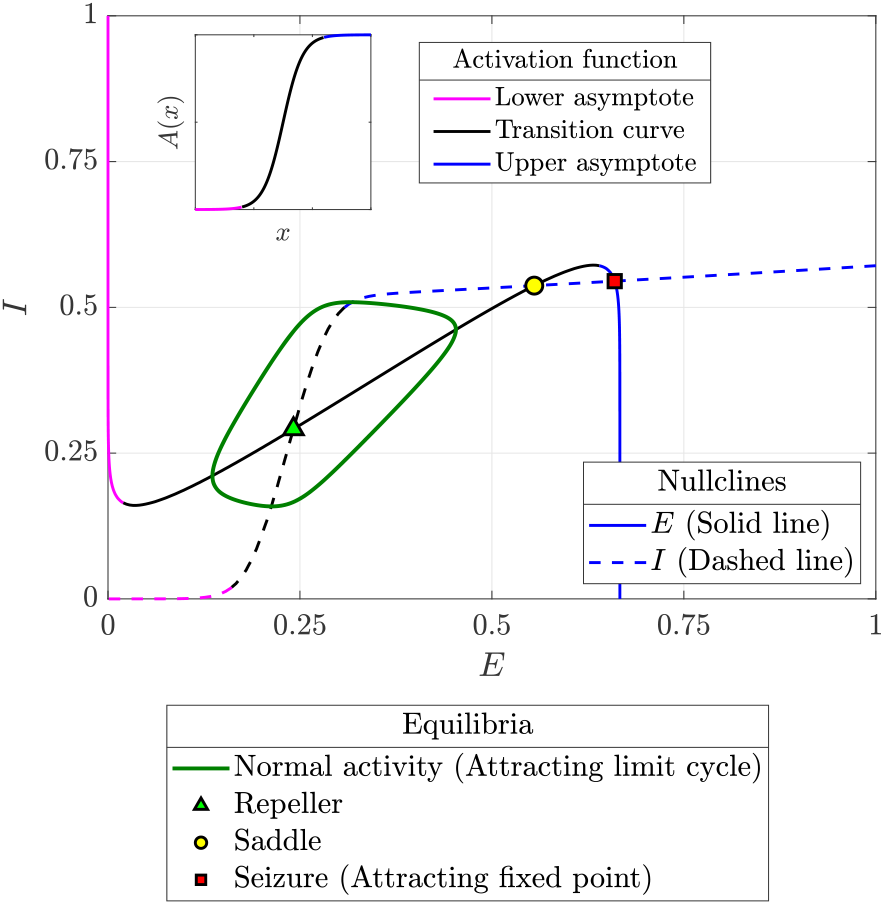
Illustrative phase portrait showing the *E* and *I* nullclines, with the different segments of the nullclines coloured to correspond with the respective segments of the activation function (inset). The different equilibria are also indicated therein: two attractors, one corresponding to normal activity and the other to seizure, a repeller, and a saddle.

We define normal activity as the attractor characterized by minimal but non-zero excitatory and inhibitory activity. This attractor arises at the first intersection of the segments of the *E* and *I* nullclines that correspond to the transition regions (plotted in black in Fig. 1) of their respective activation functions. The transition segments ensure non-zero activity, while the first intersection guarantees this activity remains minimal. Depending on system dynamics, normal activity can appear as either an attracting fixed point at this intersection or, if the fixed point is repelling, an attracting limit cycle centred around it. The latter case is illustrated in Fig. 1, where the limit cycle is shown in dark green and the repelling fixed point as a pale green triangle.

A seizure in our framework corresponds to the stable equilibrium formed at the intersection of upper asymptote segments of the *E*- and *I*-nullclines. As with normal activity, seizure can appear as either an attracting fixed point at this intersection or, if the fixed point is repelling, an attracting limit cycle centred around it. The former case is illustrated in Fig. 1, where the seizure fixed point is shown as a red square. Since the upper asymptote represents maximal activation, the seizure attractor reflects a state of runaway excitation that inhibitory mechanisms, even at full activation, cannot control.

The equilibria formed at other intersections between the *E*- and *I*-nullclines are either repellers or saddles, forming barriers that separate the two attractors of interest: normal activity and seizure.

### II.1. Baseline model

To serve as the reference point for our analyses, we establish a ‘baseline model’ with parameters configured such that a limit cycle representing normal activity is the only attractor in the system. The specific parameter values for this baseline model are listed in Table I.

**TABLE 1.** Parameters for the baseline model.

| Connection weights |  |  |  | Activation function |  |  |  |
| --- | --- | --- | --- | --- | --- | --- | --- |
| $a_{EE}$ | $a_{EI}$ | $a_{IE}$ | $a_{II}$ | $\theta_E$ | $\mu_E$ | $\theta_I$ | $\mu_I$ |
| 10 | 10 | 12 | 1 | 3 | 1.5 | 5 | 2.7 |
| Rate constants |  | External drive |  | Sustenance |  |  |  |
| $\tau_E$ | $\tau_I$ | $D_E$ | $D_I$ | $q_E$ | $q_I$ | | |
| 1 | 1 | 0.25 | 0 | 0.75 | 0.25 |  |  |

In the following sections, we examine how different excitatory and inhibitory dysfunctions modify the geometry of the nullclines, thereby shaping brain dynamics and triggering seizure onset. For each dysfunction, we begin from the baseline and progressively vary a control parameter to demonstrate how increasing dysfunction transitions the system from a single attractor corresponding to normal activity, to a bistable regime with coexisting normal and seizure states, and ultimately to seizure as the only attractor.

### II.2. Seizures arising from hyperexcitation

We model hyperexcitation as an increase in the net drive to the excitatory population (*D*_*E*_) and analyse the conditions under which it can trigger a seizure.

Figure 2 illustrates the evolution of neuronal activity over time as the control parameter *D*_*E*_ increases linearly. All parameters remain fixed at baseline values except for *D*_*E*_, which is initially held at 0.25 between *t* = 0 and *t* = 40, where the system exhibits a limit cycle corresponding to normal activity. From *t* = 40 to *t* = 90, *D*_*E*_ increases linearly from 0.25 to 2.75 and is then maintained at 2.75. Note that time *t* is a dimensionless quantity in our analyses.

**FIG. 2.**
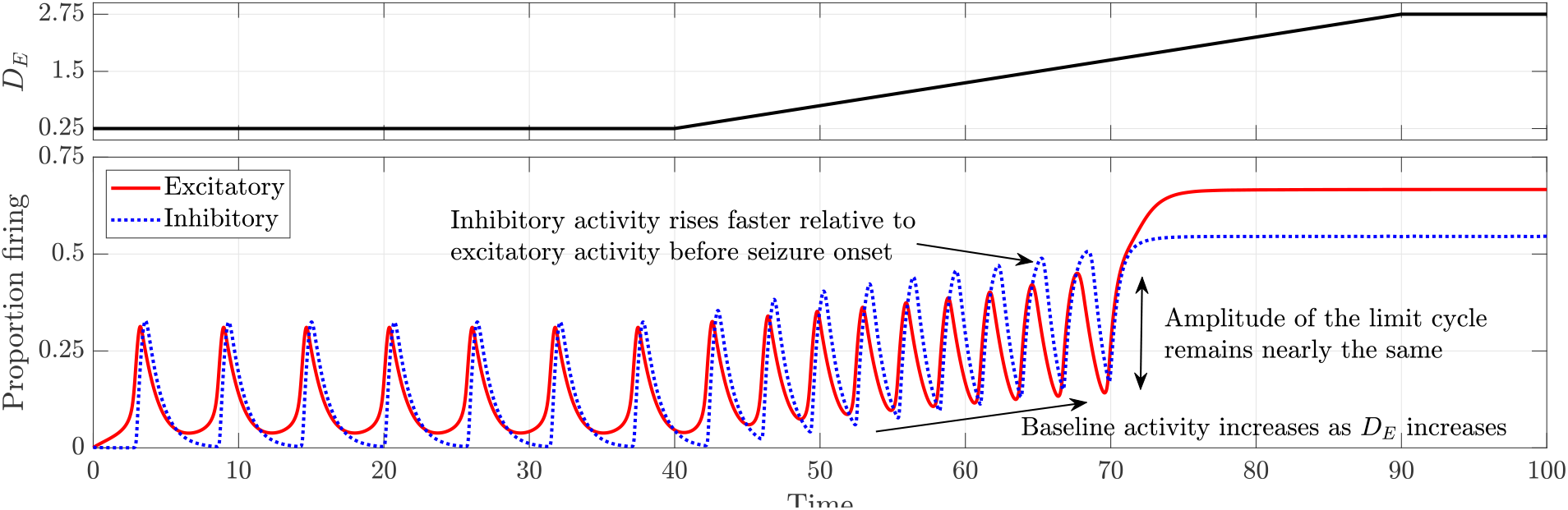
Time series depicting the transition from normal activity to seizure as the drive to the excitatory population *D*_*E*_ is increased linearly. All other parameters remain as in the baseline model.

As *D*_*E*_ increases, the limit cycle retains a nearly constant amplitude while baseline activity steadily rises. Inhibitory activity rises sharply relative to excitatory activity, reflecting inhibition’s effort to counteract the growing excitation. The transition from normal activity to seizure occurs precisely when the limit cycle describing normal activity vanishes through the saddle-homoclinic bifurcation. At the bifurcation point of *D*_*E*_ *≈* 1.7751, excitatory activity jumps suddenly, marking seizure onset. Beyond this threshold, the system remains in the seizure attractor. This transition is further illustrated through phase portraits in Fig. 3.

**FIG. 3.**
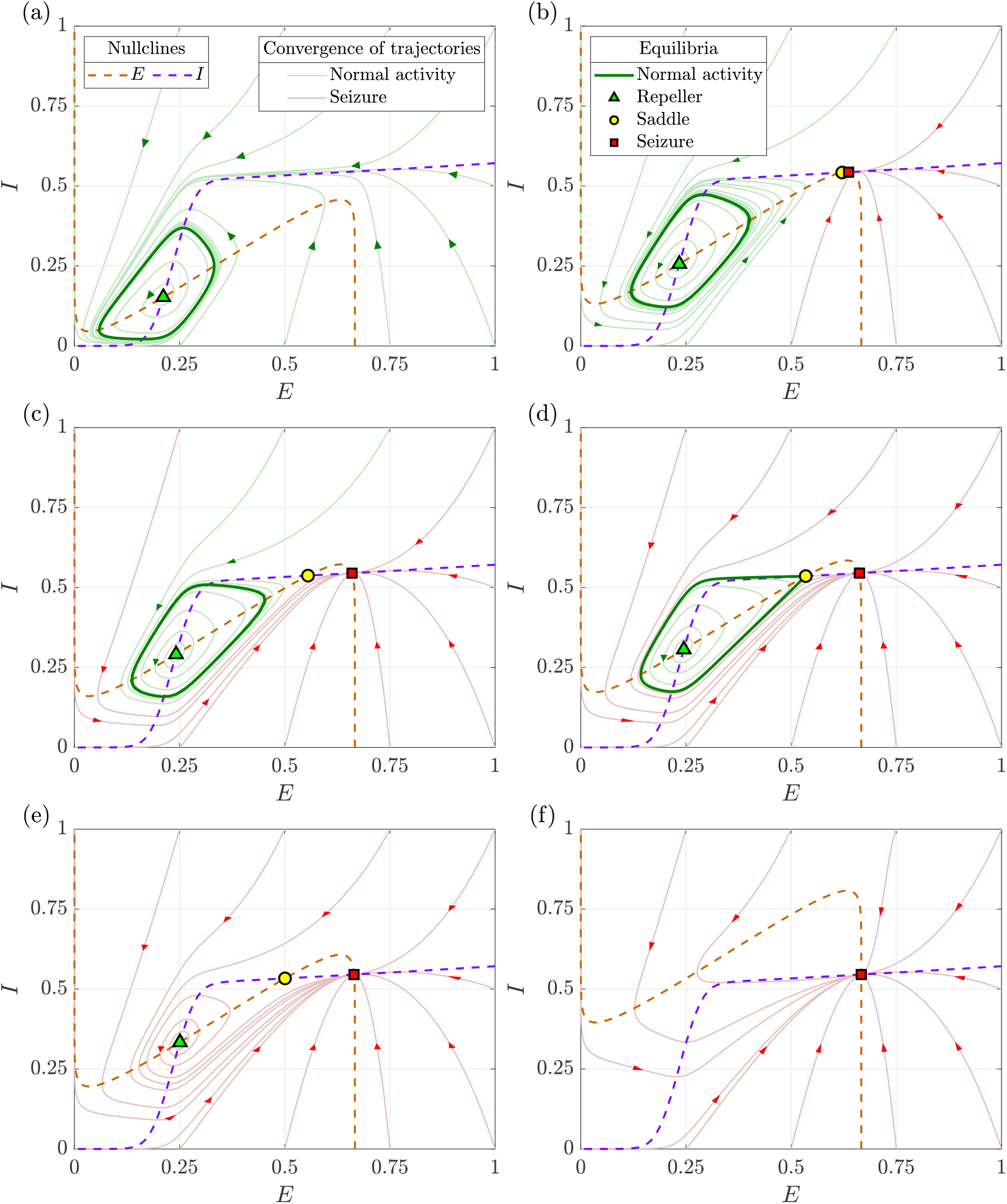
Phase portraits illustrating the change in dynamics with an increase in the drive to the excitatory population (*D*_*E*_): (a) *D*_*E*_ = 0.5: normal activity is the sole attractor, (b) *D*_*E*_ = 1.36: birth of seizure attractor and saddle through a saddle-node bifurcation, (c) *D*_*E*_ = 1.65: bistability between normal activity and seizure, (d) *D*_*E*_ = 1.7751: normal activity vanishes through a saddle-homoclinic bifurcation, (e) *D*_*E*_ = 2: seizure is the sole attractor, while the saddle and repeller persist as non-attracting equilibria, and (f) *D*_*E*_ = 4: seizure remains the sole equilibrium. All other parameters remain as in the baseline model.

Figure 3(a) illustrates the phase portrait with a low excitatory drive (*D*_*E*_ = 0.5). Here, the *E* and *I*-nullclines intersect only once and the resulting fixed point (green triangle) is repelling. A stable limit cycle (solid green curve) surrounds this point, representing normal activity. Since all trajectories converge onto this limit cycle, it is globally attracting.

Increasing *D*_*E*_ shifts the *E*-nullcline away from the *E*-axis, while the *I*-nullcline remains unchanged. At *D*_*E*_ *≈* 1.353, a second intersection between the two null-clines occurs, leading to a saddle-node bifurcation. This bifurcation creates two new fixed points: a saddle (yellow circle) and an attractor (red square), the latter corresponding to seizure. Figure 3(b) depicts the phase portrait just after this bifurcation (*D*_*E*_ = 1.36), where the emergence of the seizure attractor introduces bistability. The trajectories converging to normal activity and seizure are shown in green and red, respectively.

As *D*_*E*_ continues to increase, the limit cycle representing normal activity gradually moves closer to the saddle, as shown in Fig. 3(c). At *D*_*E*_ *≈* 1.7751, the limit cycle collides with the saddle and disappears through a saddle-homoclinic bifurcation. Figure 3(d) depicts this bifurcation point, where all trajectories originating inside the limit cycle still approach it, while those originating outside it converge onto the seizure attractor.

Further increasing *D*_*E*_ destroys bistability, making seizure the global attractor. Figure 3(e) illustrates this scenario with *D*_*E*_ = 2, where the repeller and the saddle remain as non-attracting equilibria. These two equilibria persist until *D*_*E*_ *≈* 3.4236, at which point they annihilate each other in a saddle-node bifurcation. Beyond this threshold, seizure becomes the sole equilibrium state, as depicted in Fig. 3(f).

### II.3. Seizures arising from the depletion of inhibitory neurotransmitter

Seizures can also arise from dysfunctions in inhibition. For example, when inhibitory neurons fire at high frequencies for prolonged periods, the mechanisms that sustain synaptic function, such as neurotransmitter production and vesicle recycling, may fail to keep pace with demand [31]. This can lead to depletion of inhibitory neurotransmitters like GABA at excitatory postsynaptic neurons, thereby weakening inhibitory feedback and increasing susceptibility to seizures [32, 33]. We model the propensity for inhibitory neurotransmitter depletion using the parameter *ρ ∈* [0, 1], where *ρ* = 0 indicates no depletion and *ρ* = 1 indicates maximum depletion.

Figure 4 illustrates the evolution of neuronal activity as the depletion parameter *ρ* increases linearly with time, while phase portraits depicting key stages of this transition are shown in Fig. 5. All parameters, except *ρ*, are set as in the baseline model and held constant.

**FIG. 4.**
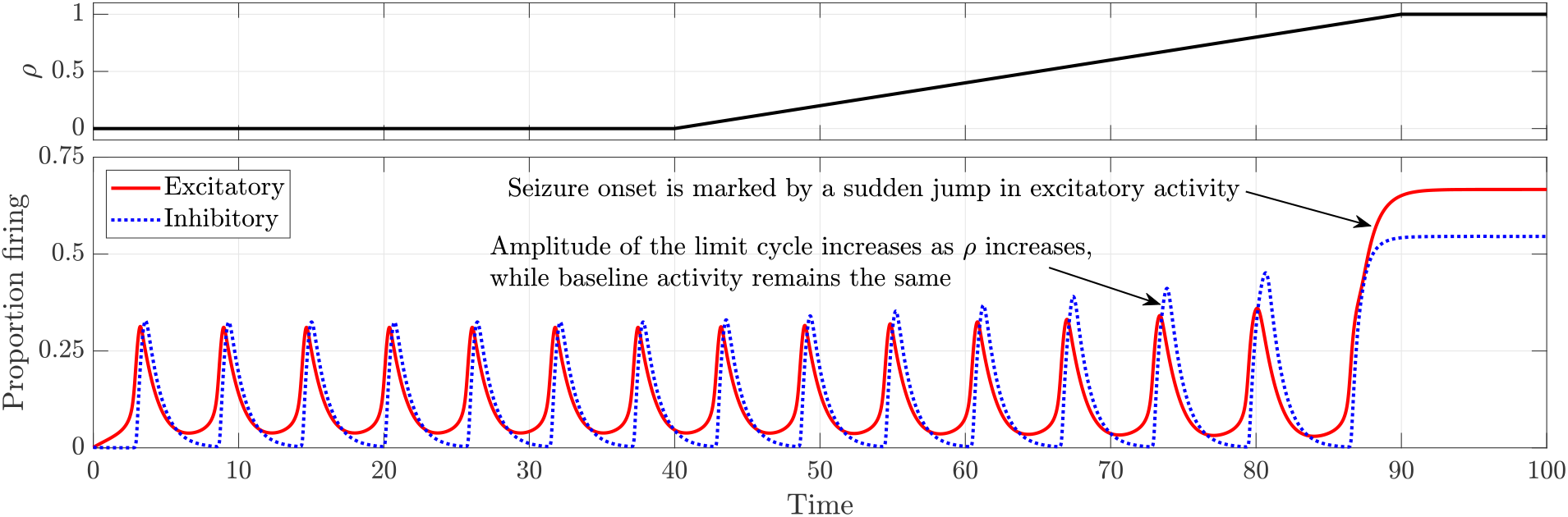
Time series depicting the transition from normal activity to seizure as the depletion parameter *ρ* is increased linearly. All other parameters remain as in the baseline model.

**FIG. 5.**
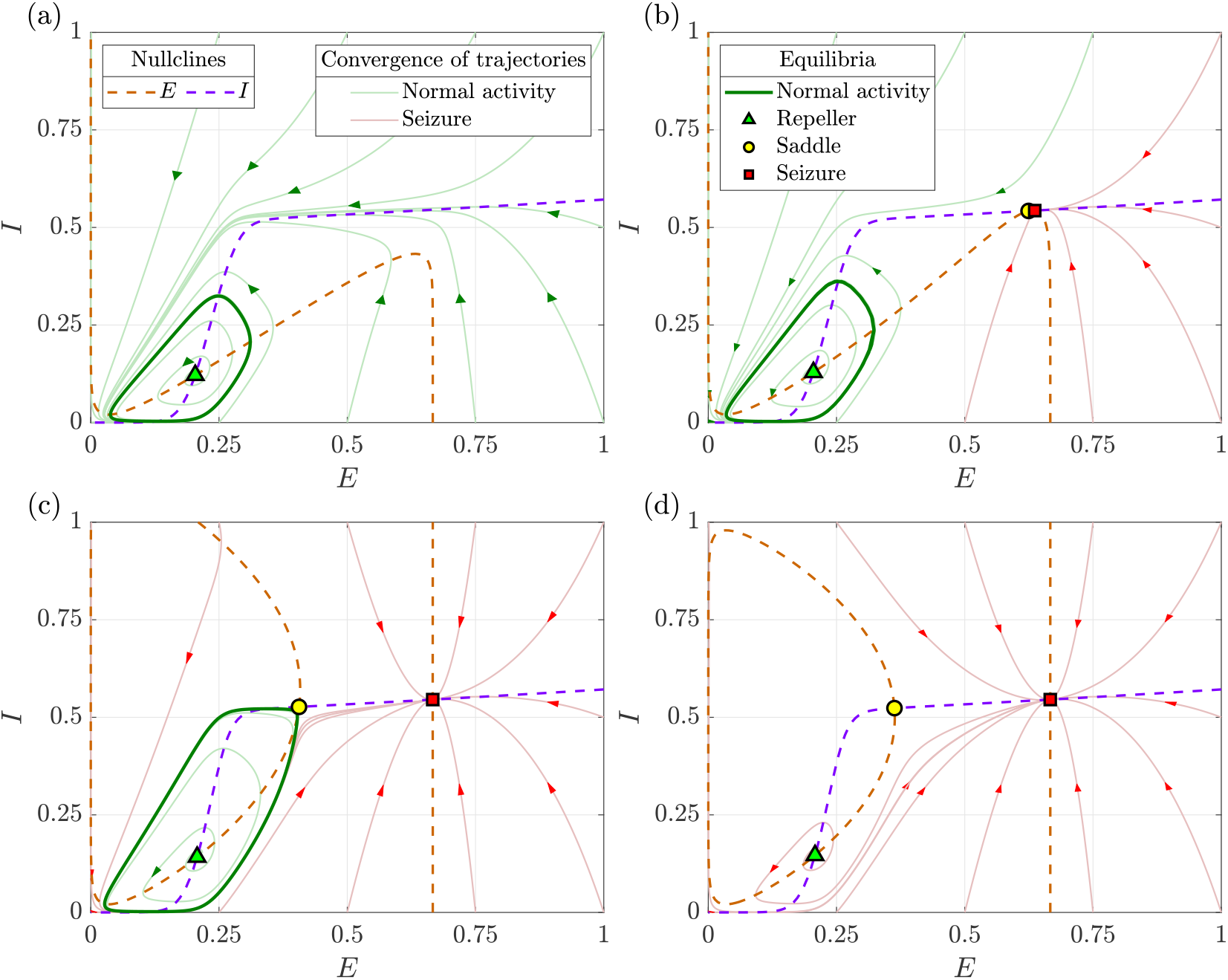
Phase portraits illustrating the change in dynamics with an increase in the depletion parameter (*ρ*): (a) *ρ* = 0: normal activity is the sole attractor, (b) *ρ* = 0.376: birth of seizure attractor and saddle through a saddle-node bifurcation leading to bistability, (c) *ρ* = 0.874: normal activity vanishes through a saddle-homoclinic bifurcation, and (d) *ρ* = 1: seizure remains the sole attractor. All other parameters remain as in the baseline model.

At *ρ* = 0, the system exhibits a globally attracting limit cycle corresponding to normal activity as shown in Fig. 5(a). As *ρ* increases from *t* = 40, the time series in Fig. 4 shows that the limit cycle slowly grows in amplitude, while the baseline activity remains unchanged. This contrasts with the case of increased drive to the excitatory population, where baseline activity rises while the amplitude of the limit cycle remains constant.

The phase portraits in Fig. 5 show that increasing *ρ* primarily affects the *E*-nullcline, shifting it away from the *E*-axis. At *ρ ≈* 0.3744, *E*- and *I*-nullclines intersect again, giving rise to a saddle and the seizure attractor through a saddle-node bifurcation. Figure 5(b) depicts the phase portrait just after this bifurcation, illustrating bistability. Notably, the time series does not immediately reflect this transition from monostability to bistability.

As *ρ* increases further, the limit cycle continues to expand until it eventually collides with the saddle and disappears through a saddle-homoclinic bifurcation at *ρ ≈* 0.874 as shown in Fig. 5(c). This destroys bistability, leaving seizure as the sole attractor: for *ρ* > 0.874, all trajectories converge onto the seizure attractor, as illustrated in Fig. 5(d). In the time series, this saddle-homoclinic bifurcation corresponds to seizure onset, characterized by a sudden jump in excitatory activity as the trajectory transitions from the limit cycle to the seizure attractor.

### II.4. Seizures arising from the depolarising effect of GABAergic neurotransmission

The inhibitory neurotransmitter gamma-aminobutyric acid (GABA) exerts its effects predominantly through the activation of GABA_*A*_ receptors on neurons. Under normal conditions, activation of these receptors opens chloride channels, resulting in chloride influx and neuronal hyperpolarization, provided that the chloride equilibrium potential is more negative than the resting membrane potential [13]. However, in developmental stages or pathological conditions such as epilepsy, alterations in chloride transporter expression can disrupt this balance. Specifically, an imbalance between chloride influx via the sodium–potassium–chloride co-transporter (NKCC1) and efflux via the potassium–chloride co-transporter (KCC2) can lead to persistent intracellular chloride accumulation [34]. We model this transporter imbalance using the parameter *κ*, where *κ* = 0 represents intact homeostasis and positive values indicate impairment, with higher values reflecting greater dysfunction.

When intracellular chloride levels become sufficiently elevated, the chloride reversal potential shifts toward more depolarized values, causing a switch in GABA_*A*_ receptor activation from chloride influx to efflux. This reversal of GABA’s effect from inhibition to excitation undermines inhibitory restraint and can promote seizures [35, 36]. Accordingly, we explore the evolution of neuronal activity by gradually increasing the chloride accumulation parameter *κ* over time. The sensitivity parameter quantifying the effect of depolarising GABA on postsynaptic neurons, *a*_*PI*_, is held constant at 5. All other parameters are set as in the baseline model and held constant. The results, shown in Fig. 6, closely resemble those observed in the case of inhibitory neurotransmitter depletion, with the transition from normal activity to seizure occurring via a saddle-homoclinic bifurcation at *κ ≈* 1.61714.

**FIG. 6.**
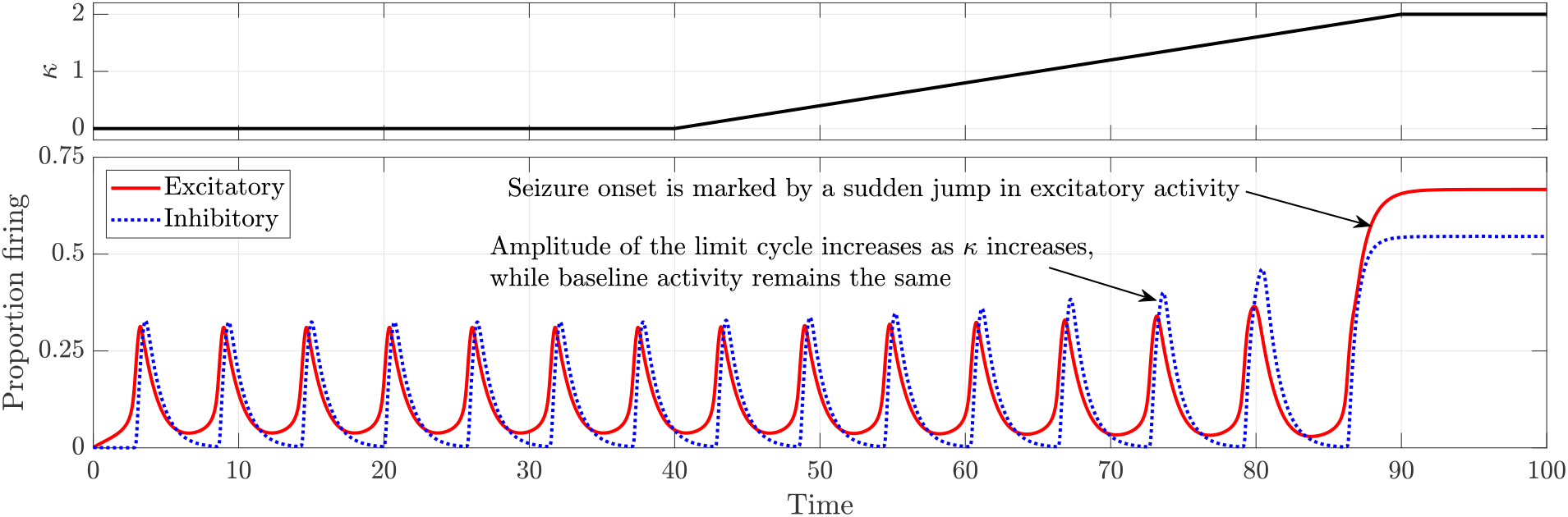
Time series depicting the transition from normal activity to seizure as the chloride accumulation parameter *κ* is increased linearly. All other parameters remain as in the baseline model.

### II.5. Interventions for seizure termination are influenced by the underlying dysfunction

Having analysed how various dysfunctions contribute to seizure onset, we now turn to their implications for interventions aimed at seizure termination. In our model, seizure termination is defined as the disappearance of the seizure attractor via a saddle-node bifurcation, rather than simply restoring bistability. In a bistable regime, noise can induce transitions both into and out of the seizure attractor; thus, eliminating the seizure attractor is essential to prevent immediate seizure recurrence.

Since benzodiazepines, the first-line treatment, terminate seizures by enhancing GABAergic inhibition, we begin by investigating the effects of GABAergic enhancement on neuronal dynamics within our model. We represent GABAergic enhancement by introducing a factor, *σ*_GABA_, which amplifies the effect of inhibition in the argument of the excitatory activation function. To illustrate this, we first consider seizures driven by hyperexcitation. All parameters are set as in the baseline model, except for the drive to the excitatory population, which is set to *D*_*E*_ = 3 to ensure seizure remains the sole attractor.

Figure 7 illustrates the effect of three levels of GABAergic enhancement on excitatory activity (*E*). Inhibitory activity is omitted for clarity, as excitatory activity alone is sufficient to determine whether the brain is in a seizure or normal state. Between *t* = 0 and *t* = 20, GABAergic enhancement is held at the baseline level of *σ*_GABA_ = 1 and the system remains in a seizure state.

**FIG. 7.**
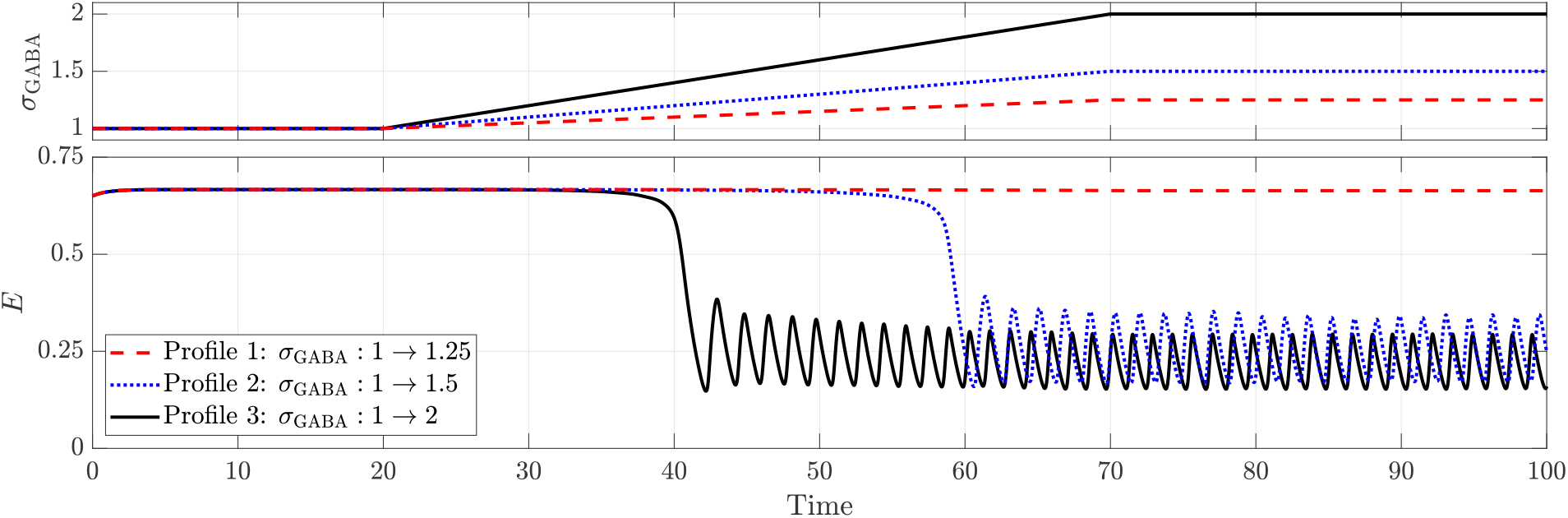
Effect of GABAergic enhancement in terminating seizures caused due to hyperexcitation (*D*_*E*_ = 3). All other parameters remain as in the baseline model.

Between *t* = 20 and *t* = 70, *σ*_GABA_ is increased linearly to three different levels, 1.25, 1.5 and 2, and held at these levels thereafter.

When *σ*_GABA_ peaks at 1.25, seizure termination is unsuccessful. However, higher levels (*σ*_GABA_ = 1.5 and 2) successfully terminate the seizure, restoring normal activity characterized by a limit cycle. Additionally, comparing *σ*_GABA_ = 1.5 and 2 reveals that stronger GABAergic enhancement results in faster seizure termination and a lower proportion of neurons firing during normal activity.

A similar effect is observed when GABAergic enhancement is employed to terminate seizures caused by the depletion of inhibitory neurotransmitter as shown in Fig. 8, with two key differences. First, successful termination occurs at higher levels of GABAergic enhancement in the depletion case compared to seizures driven by hyperexcitation. Second, in the depletion case, the proportion of neurons firing during normal activity remain largely unchanged regardless of the level of GABAergic enhancement.

**FIG. 8.**
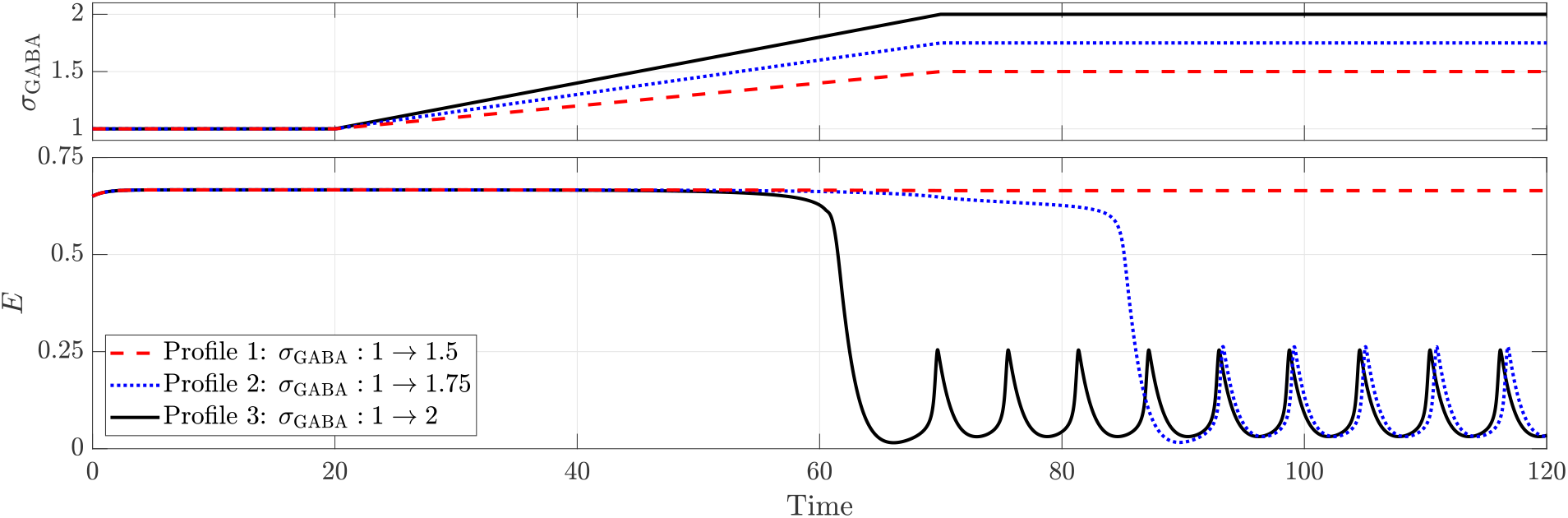
Effect of GABAergic enhancement in terminating seizures caused due to the depletion of inhibitory neurotransmitter (*ρ* = 1). All other parameters remain as in the baseline model.

On the other hand, when GABAergic neurotransmission becomes depolarising, GABAergic enhancement fails to terminate the seizure as shown in Fig. 9. This failure occurs because GABA’s role switches from inhibitory to excitatory, so enhancing its effect no longer suppresses seizure activity.

**FIG. 9.**
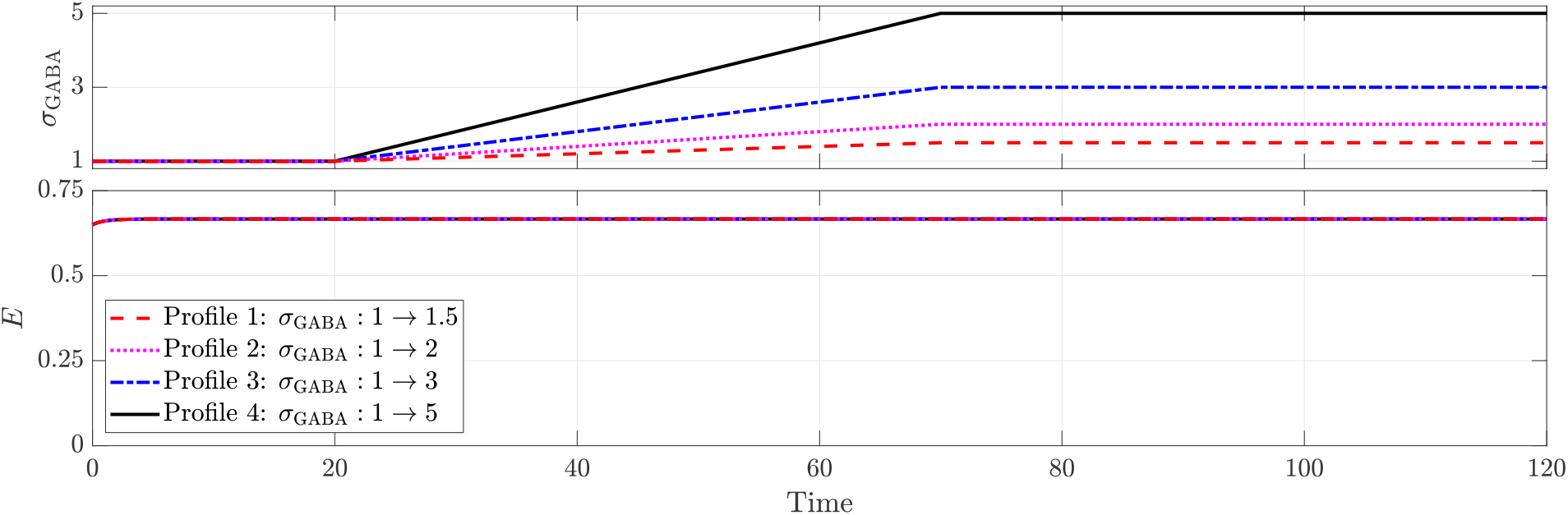
GABAergic enhancement fails to terminate seizures when GABAergic neurotransmission is depolarising (*κ* = 1.8). All other parameters remain as in the baseline model.

To terminate such benzodiazepine-refractory seizures, second-line treatments such as levetiracetam and phenytoin, which selectively antagonize rhythmic firing, are recommended [15]. In our framework, the sustenance term encodes factors responsible for persistent firing. Consequently, to model the selective suppression of rhythmic activity, we introduce a term *σ*_RS_ that counter-acts sustenance by modifying Eqs. (7) and (8), replacing *q*_*E*_ and *q*_*I*_ with (*q*_*E*_ *− σ*_RS_) and (*q*_*I*_ *− σ*_RS_) respectively.

Figure 10 illustrates the effect of varying levels of rhythmic suppression on excitatory activity (*E*). From *t* = 0 to *t* = 20, there is no rhythmic suppression (*σ*_RS_ = 0), and the system remains in a seizure state. Between *t* = 20 and *t* = 70, *σ*_RS_ is increased linearly to four different levels, 0.5, 1, 1.5, and 2, after which it is held constant.

**FIG. 10.**
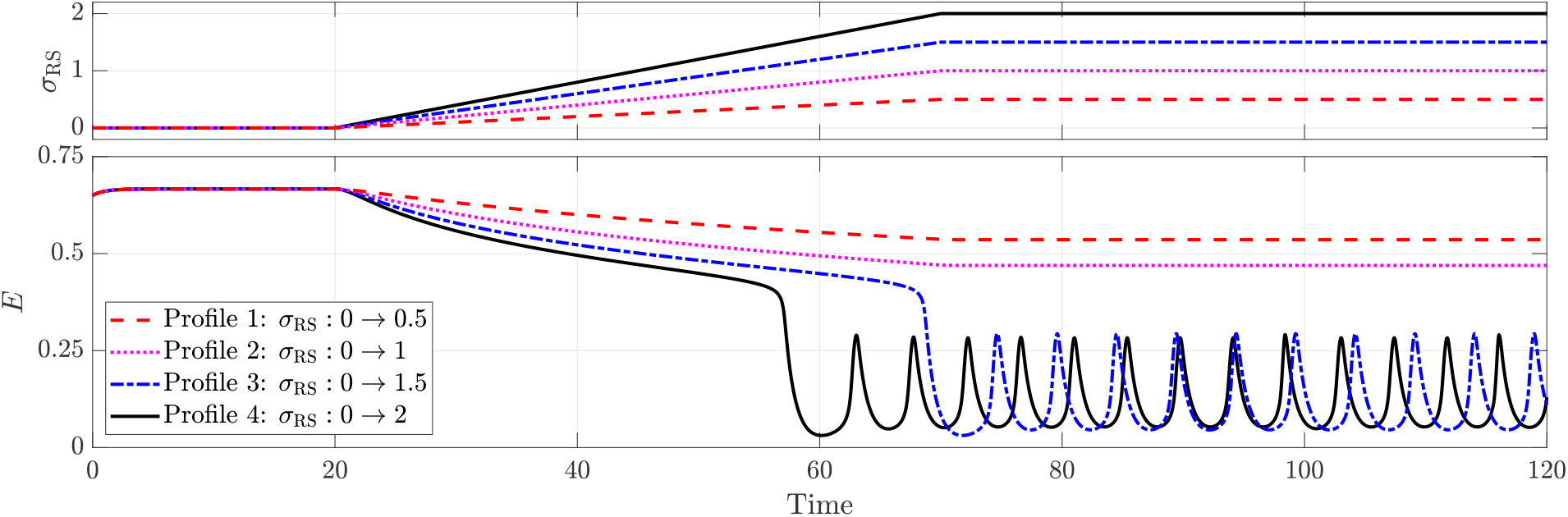
Effect of rhythmic suppression (*σ*_RS_) in terminating seizures driven by depolarising GABAergic neurotransmission (*κ* = 1.8). All other parameters remain as in the baseline model.

Seizure termination occurs when *σ*_RS_ reaches 1.5 or higher, with higher levels leading to faster termination. While lower levels of rhythmic suppression do not fully stop the seizure, they still reduce neuronal activity during seizures — an effect not observed with GABAergic enhancement. Moreover, increasing *σ*_RS_ results in a greater suppression of activity, and for levels sufficient to achieve termination, higher doses lead to a slight increase in the frequency of the limit cycle representing normal activity.

To understand why rhythmic suppression can terminate seizures when GABAergic enhancement fails, we compare their effects on the *E*- and *I*-nullclines under conditions of depolarizing GABAergic neurotransmission (Fig. 11). Rhythmic suppression, by counteracting the sustenance term, shifts the upper asymptote segments of both nullclines that underpin the seizure attractor and alters the position of the seizure attractor in phase space. Specifically, it moves the *E*-nullcline toward the *I*-axis and the *I*-nullcline toward the *E*-axis, thereby reducing the level of activity during seizure. This progressive shift ultimately leads to the collision and mutual annihilation of the seizure attractor and the adjacent saddle point, resulting in seizure termination. In contrast, GABAergic enhancement via benzodiazepines affects only the excitatory activation function and does not shift the seizure attractor. Consequently, it neither reduces neuronal activity during seizures nor achieves termination. These findings underscore the importance of aligning therapeutic strategies with the specific pathophysiological mechanisms for effective seizure termination.

**FIG. 11.**
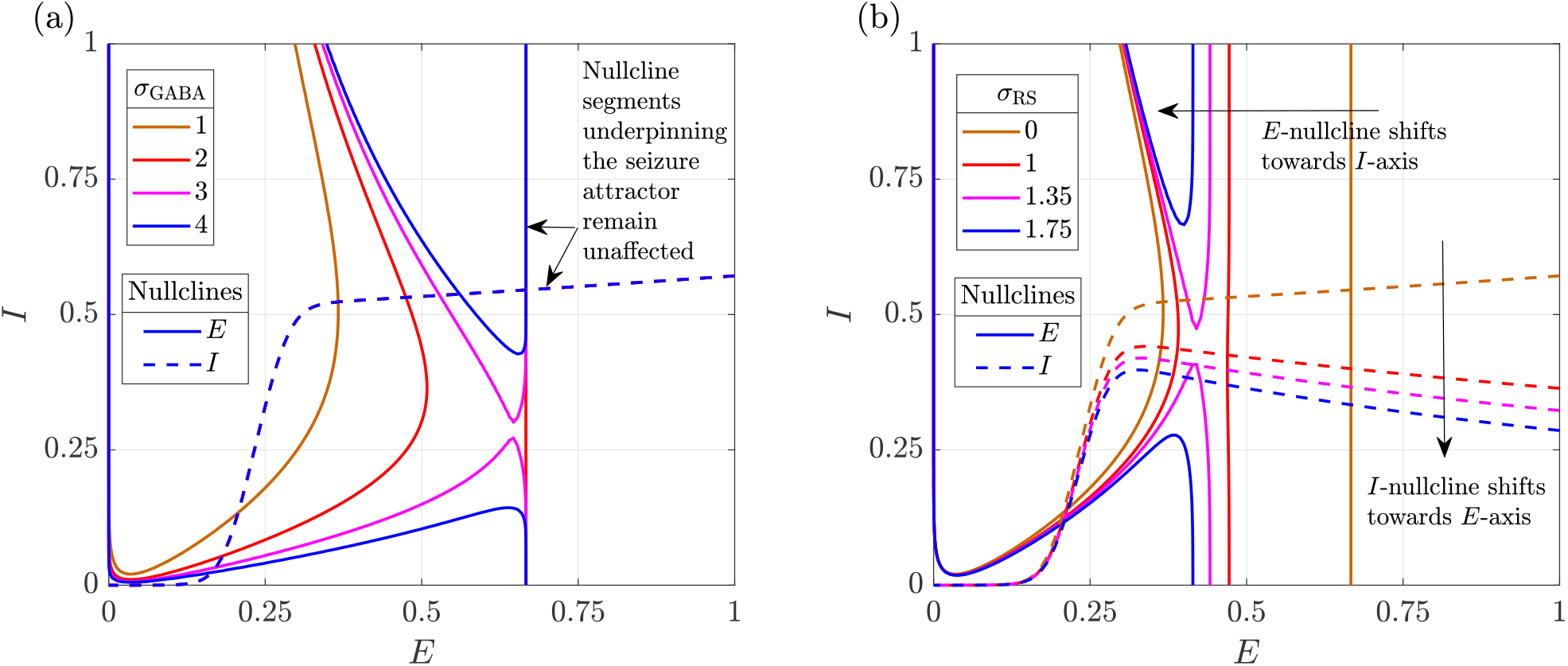
Influence of (a) GABAergic enhancement and (b) rhythmic suppression on the *E* and *I* nullclines when GABAergic neurotransmission is depolarising (*κ* = 1.8). All other parameters remain as in the baseline model.

## III. DISCUSSION

We now turn to a discussion of the key features of our model, its limitations, and potential avenues for future extensions.

A key innovation in our framework is the introduction of a second-order interaction term, which quantifies mechanisms that promote or discourage persistent neuronal firing. We begin by modelling mechanisms that promote sustained activity, referring to this contribution as the sustenance term. Several biophysical processes can underlie such positive feedback. For instance, the activation of N-methyl-D-aspartate (NMDA) receptors is known to drive persistent firing of excitatory cells and is dependent upon sufficient depolarization [37]. Changes in NMDA receptor abundance or subunit composition may further amplify this effect [38, 39] and in our framework, higher levels of sustenance can phenomenologically reflect such amplification. The sustenance term can also compactly encode the extent of abnormal recurrent excitatory connections that can reinforce pathological activity patterns [40].

To capture the influence of therapeutic interventions that suppress pathological rhythmic firing such as phenytoin and levetiracetam, we extend the model by incorporating terms that counteract the effects of sustenance. Although phenytoin and levetiracetam act through distinct mechanisms, both effectively suppress abnormal firing in epileptic circuits while sparing normal electrophysiological function [41, 42]. Our model phenomenologically reproduces this suppression, and the results support their efficacy as alternative therapies for benzodiazepine-refractory seizures, consistent with clinical observations [43, 44].

Previous approaches have modelled suppression at high excitatory input by replacing the sigmoidal activation function with a Gaussian [45]. However, accurately reproducing partial suppression rather than complete silencing requires a finely tuned combination of a Gaussian and a sigmoid, which increases model complexity and reduces interpretability. In contrast, our sustenance term offers an elegant and biologically grounded formulation for capturing persistent activity and its modulation, striking a balance between model fidelity and complexity.

In the original Wilson-Cowan model [30], normal activity is typically represented by the trivial fixed point (0, 0). In contrast, normal activity in our framework is nontrivial, better reflecting physiological observations that a small but non-zero fraction of excitatory and inhibitory neurons remain active in a healthy brain. Furthermore, the seizure attractor corresponds to uncontrolled excitation that persists despite maximal inhibitory activation. While earlier studies on neuronal ensembles have explored non-trivial, oscillatory equilibria to represent normal activity [46, 47], our work offers a formal definition within the Wilson-Cowan framework: normal activity is defined as the stable attractor arising from the first intersection of the transition segments of the *E*- and *I*-nullclines whereas whereas seizure corresponds to the stable attractor formed by the intersection of the upper asymptotic segments. This formalism enables a precise and physiologically interpretable classification of the dynamic regimes in epilepsy.

For most individuals with chronic epilepsy, seizures are rare events, accounting for less than 1% of total brain activity. This suggests that normal activity serves as the predominant attractor, while seizures result from occasional, activity-dependent imbalances in excitatory and inhibitory neurotransmission [48]. To reflect this, we have included an activity-dependent contribution whilst modelling dysfunctions. Specifically, the model for inhibitory neurotransmitter depletion assumes that the level of depletion is proportional to the average inhibitory activity over a preceding time window. Similarly, the model for depolarising GABAergic neurotransmission reflects the dependence of neuronal depolarization and GABA_*A*_ activation on ongoing excitatory and inhibitory activity, respectively. While modelling hyperexcitation, the increased excitatory drive *D*_*E*_ is not constructed to be explicitly activity-dependent; however, it implicitly captures elevated excitatory input from neighbouring neuronal populations.

Our analyses reveal that pathological perturbations, whether excitatory or inhibitory, primarily impact the *E*-nullcline. Regardless of the specific dysfunction, as the control parameter varies, the seizure attractor emerges via a saddle-node bifurcation, while normal activity disappears through a saddle-homoclinic bifurcation. The parameter space between these bifurcations corresponds to a bistable regime, where both normal activity and seizure coexist. Although seizure onset occurs at the saddle-homoclinic bifurcation in a noise-free system, real-world neural activity is inherently noisy. Consequently, the transition to seizure can occur at any point after the seizure attractor appears.

Furthermore, our simulations consistently show that, across different dysfunctions, inhibitory activity rises sharply relative to excitation just before seizure onset. The onset itself is then marked by a sudden surge in excitatory activity. This dynamic aligns with physiological observations that inhibitory activity often intensifies prior to seizure onset [49].

While all dysfunctions primarily affect the *E*-nullcline, the way each one alters its geometry is unique, and these changes are reflected indirectly in the time series through the behaviour of the limit cycle. Specifically, inhibitory dysfunctions cause the amplitude of the limit cycle corresponding to normal activity to increase before seizure onset, whereas hyperexcitation raises baseline activity and increases the frequency of the limit cycle without altering its amplitude.

Previous work has demonstrated that features such as EEG amplitude, inter-spike intervals, and the presence of a DC shift can be used to classify seizure onset and offset into various co-dimension one bifurcations [50, 51]. Similar techniques can be employed to identify these dysfunctions through specific EEG features that reflect the unique changes in limit cycle behaviour and guide targeted interventions.

In terms of interventions, GABAergic enhancement by benzodiazepines primarily influences the geometry of the *E*-nullcline, whereas suppression of rhythmic firing exerts a substantial influence on both the *E*- and *I*-nullclines. When seizures are driven by hyperexcitation or depletion of inhibitory neurotransmitters, shifting the *E*-nullcline alone is sufficient for seizure termination — an effect successfully achieved through GABAergic enhancement. However, when GABAergic neurotransmission becomes depolarizing, enhancing GABAergic inhibition no longer shifts the *E*-nullcline sufficiently to eliminate its intersection with the *I*-nullcline that generates the seizure attractor. Instead, it may restore bistability between seizure and normal activity. Ultimately, as the seizure attractor persists despite the intervention, if the system begins in the seizure state, it remains trapped there.

In contrast, suppression of rhythmic firing directly perturbs the seizure attractor by reducing the proportion of neurons firing at maximal activation. By shifting the upper asymptote segments of the *E*- and *I*-nullclines, it brings the seizure attractor and the adjacent saddle point together until they collide and annihilate through a saddle-node bifurcation. This eliminates the seizure attractor and restores normal activity.

These findings highlight the need for a systematic approach to treatment selection. While control-theoretic approaches have been explored for optimizing brain stimulation in seizure termination [52, 53], similar systematic studies for drug-based interventions remain limited. Our framework provides a foundation for addressing this gap, enabling the optimization of drug type, dosage, and timing by minimizing a cost function over the parameter space.

Our current model does not account for shunting inhibition, a mechanism by which inhibitory neurons reduce excitability by increasing membrane conductance and diverting electrical currents [54, 55]. Shunting inhibition is particularly relevant when GABAergic currents are depolarizing, as the resting membrane potential remains slightly above the chloride reversal potential. In this scenario, even if chloride efflux occurs, the increased conductance allows inhibitory shunting to counteract excitation, preserving the net inhibitory effect of GABA [56]. Future extensions of our model will feature shunting inhibition through a hybrid formulation [57, 58].

Since seizures arise from abnormal synchronous neuronal firing, incorporating synchronization into our model is a natural next step. Recent extensions of the Wilson-Cowan model have integrated synchronization [59], and similar approaches can enhance our framework. Given that epilepsy is fundamentally a disorder of cortical network disorganization [60], extending the model to a network of interconnected excitatory and inhibitory populations is essential. Future work will focus on expanding our framework to multiple populations and assessing how insights from the two-population model translate to network-level dynamics.

## IV. METHODS

We begin with the original Wilson–Cowan equations, which describe the dynamics of interacting excitatory and inhibitory neuronal populations, and progressively incorporate additional features to capture mechanisms relevant to epilepsy. The Wilson–Cowan model is given by [30],

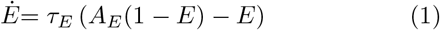

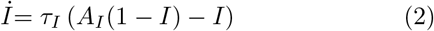

with dynamics restricted to Ω = [0, 1] *×* [0, 1]. The state variables *E* and *I*, respectively, denote the proportion of excitatory and inhibitory neurons engaged in firing at any given time *t*. The parameters *τ*_*E*_ and *τ*_*I*_ denote the rate constants for the excitatory and inhibitory populations, respectively. The activation functions *A*_*E*_ = *f* (*x*_*E*_) and *A*_*I*_ = *g*(*x*_*I*_) govern the recruitment into firing for excitatory and inhibitory populations, respectively. The arguments, *x*_*E*_ and *x*_*I*_, encode how quiescent neurons perceive their input: a weighted sum of excitatory *E* and inhibitory *I* activity, and a net drive,

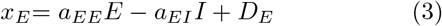

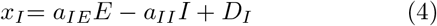

where, *D*_*E*_ and *D*_*I*_ represent the net drive for the excitatory and inhibitory populations, respectively, and *a*_*ii*_ represent the relative weight of connections within and between the two populations.

Given that a higher net excitatory activity can promote recruitment, activation functions are typically considered to be monotonically increasing functions, such as a sigmoid [61]. We choose,

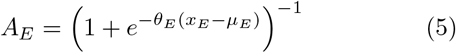

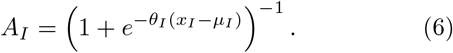

Both activation functions have a lower asymptote at zero and an upper asymptote at one. The parameter *µ*_*i*_ specifies the value of *x*_*i*_ at which the activation function reaches its median value of 0.5 and the parameter *θ*_*i*_ controls the slope of the sigmoid. Different choices for activation functions may lead to variations in the specific dynamics, but the stability of the equilibria of interest would remain unchanged [30].

A defining feature of epileptic seizures is persistent neuronal firing. When a large fraction of excitatory neurons become active, this heightened activity can sustain pathological firing through positive feedback. To model this mechanism, we introduce a term referred to as sustenance by replacing the original linear decay in Eqs. (1) and (2) with a second-order decay:

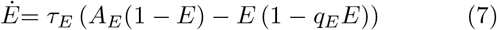

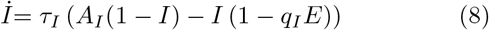

with dynamics restricted to Ω. The level of sustenance within the excitatory and inhibitory populations is quantified by the parameters *q*_*E*_ *∈* [0, 1] and *q*_*I*_ *∈* [0, 1] respectively. Equations (7) and (8) together represent the equations governing the interaction between one excitatory and one inhibitory neuronal population.

Accordingly, the excitatory nullcline is given by,

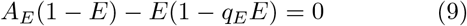

and the inhibitory nullcline is given by,

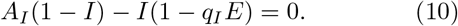

### IV.1. Modelling dysfunctions

#### IV.1.1. Depletion of inhibitory neurotransmitter

We model the depletion of inhibitory neurotransmitters as an activity-dependent process: the level of inhibitory neurotransmitter available at time *t* is influenced by the average inhibitory activity over a preceding time window Δ*t*. This depletion reduces the effective inhibition experienced by postsynaptic neurons, denoted as *I*^eff^, which we define as,

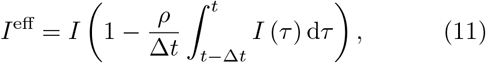

where *ρ ∈* [0, 1] is a constant that governs the level of depletion in inhibitory neurotransmitters.

Applying Taylor’s theorem to expand the integral in Eq. (11) about Δ*t* = 0 yields,

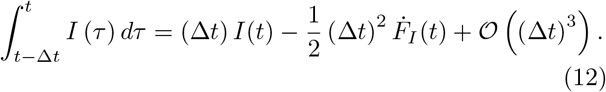

Taking the first-order approximation of the above expansion and substituting into Eq. (11) yields,

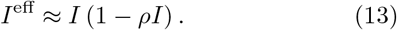

To incorporate inhibitory neurotransmitter depletion into our model, we replace *I* with *I*^eff^ in Eqs. (3) and (4), encoding the reduced inhibition felt by quiescent excitatory and inhibitory populations, while keeping the rest of the framework unchanged.

#### IV.1.2. Depolarising GABAergic neurotransmission

Depolarizing GABAergic neurotransmission occurs when postsynaptic neurons accumulate chloride ions due to disrupted chloride homeostasis. Two key factors contribute to this disruption: (1) dysfunction of chloride transporters and (2) activity-dependent chloride influx.

To model activity-dependent chloride influx, we note that excitatory activity depolarizes quiescent neurons, and when GABA_*A*_ receptors are activated, chloride influx follows. If chloride co-transporters fail to regulate this influx effectively, it can cause chloride accumulation. We model this transporter imbalance using the parameter *κ*, where *κ* = 0 represents intact homeostasis and positive values indicate impairment, with higher values reflecting greater dysfunction.

Accordingly, we define the fraction of quiescent excitatory neurons with excessive chloride accumulation as: *p* = *κEI*. This expression captures the interplay between neuronal depolarization (*E*), GABA_*A*_ activation (*I*), and transporter dysfunction (*κ*) on promoting chloride accumulation.

With this definition, we subdivide the quiescent excitatory population into two subpopulations. The first subpopulation, comprising the fraction *p*, consists of neurons with high chloride accumulation, which perceive GABAergic input as excitatory:

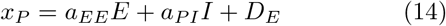

where *a*_*PI*_ represents the sensitivity of these neurons to the excitatory action of GABA. The second subpopulation, comprising the fraction 1 *− p*, consists of neurons for which GABAergic input remains inhibitory. Their perception of inputs remain identical to those of the standard excitatory population, as described by Eq. (3).

Consequently, the resultant perception by the quiescent excitatory population, combining the two subpopulations can be expressed as,

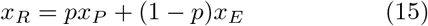

To incorporate depolarising effect of GABA into our model, we replace *x*_*E*_ with *x*_*R*_ in the activation function governing the excitatory population (Eq. (5)), while keeping the rest of the framework unchanged.

### IV.2. Modelling interventions

We model GABAergic enhancement by introducing a factor, *σ*_GABA_, to amplify the effect of inhibition, in the argument of the excitatory activation function,

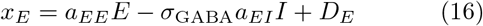

Equation (16) is used in lieu of Eq. (3) in the excitatory activation function to simulate GABAergic enhancement, while the rest of the framework remains unchanged.

To model the suppression of rhythmic firing, we introduce a term *σ*_RS_ that counteracts sustenance by modifying Eqs. (7) and (8), replacing *q*_*E*_ and *q*_*I*_ with (*q*_*E*_ *− σ*_RS_) and (*q*_*I*_ *− σ*_RS_) respectively.

### IV.3. Numerical simulation

All analyses have been carried out in MATLAB. The nullclines are plotted using the function *fimplicit* and the fixed points are obtained using the function *solve*. Limit cycles and other trajectories were simulated using the function *ode23t*.

## V. DATA AVAILABILITY

This study is based exclusively on simulated data generated specifically for the purposes of analysis and manuscript preparation. No real-world data were used at any stage. The complete simulated dataset, along with the code used for its generation, is available from the corresponding author upon reasonable request.

